# Frontoparietal, cerebellum network codes for accurate intention prediction in altered perceptual conditions

**DOI:** 10.1101/2020.11.26.399782

**Authors:** L. Ceravolo, S. Schaerlaeken, S. Frühholz, D. Glowinski, D. Grandjean

## Abstract

Integrating and predicting intentions and actions of others are crucial components of social interactions, but the behavioral and neural underpinnings of such mechanisms in altered perceptual conditions remain poorly understood. We demonstrated that expertise was necessary to successfully understand and evaluate communicative intent in spatially and temporally altered visual representations of music plays, recruiting frontoparietal regions and several sub-areas of the cerebellum. Functional connectivity between these brain areas revealed widespread organization, especially in the cerebellum. This network may be essential to assess communicative intent in ambiguous or complex visual scenes.

---

Human ability to coordinate with others is a key evolutionary skill, allowing us to achieve tasks that could not be managed individually. Beyond vocal and semantic communication (Kotz and Schwartze 2010), such mechanism relies on fine-tuned non-verbal expressive behaviors that must be able to communicate one’s intent reliably and efficiently. Intention therefore involves the whole body as a mean of communication, especially focusing on the upper body actions and movement dynamics (Andersen and Cui 2009). It requires both parties to share similar representations at different levels (e.g., common goal or intermediate steps to achieve a final goal), to predict shared outcomes and to integrate the predicted consequences of our own actions as well as others’ (Sebanz and Knoblich 2009). Hence, it also requires to pay attention to one’s own intentions and to be able to predict and anticipate movement generation (Lau, Rogers et al. 2004). However, the neural processes underlying such communication flow are not yet fully understood. Currently, the literature points at the use of an internal forward model that optimizes one’s motor control by comparing the actual and predicted sensory consequence of movements (Wolpert, Doya et al. 2003). By extension, such model might also be used to predict others’ actions in a social interaction based on one’s own action representations (Wolpert, Doya et al. 2003). Such abilities would be supported by several interacting brain networks that are active both during action generation and during observation of others’ actions (Rizzolatti, Fadiga et al. 1996). The frontoparietal network including the inferior parietal lobule (IPL), have been associated with observation of individual movements (in both monkeys and humans), shared visual attention, and motor intention recognition (Rizzolatti and Sinigaglia 2010). Attention to one’s own intentions and actions also recruited the frontoparietal network, especially the prefrontal cortex exhibiting a strong functional coupling with the premotor cortex (Lau, Rogers et al. 2004). The posterior parietal cortex, on the other hand, was repeatedly observed in situations pertaining to motor intention and imagery (Jeannerod 1994), while the lateral parietal cortex, such as the intraparietal sulcus (IPS) and IPL, were linked more directly and precisely to attention and intention itself (Lau, Rogers et al. 2004, Desmurget and Sirigu 2012, Eskenazi, Rueschemeyer et al. 2015). While such simulation system has been previously described as a mirror neuron system in monkeys based on single cell recordings, this mirror neuron system is difficult to observe properly in humans. Recognition of intentions is notably impacted by contextual information and prior knowledge (Brass, Schmitt et al. 2007). It is subject to information availability and processing (full body vision vs. occluded vision), context richness (familiar vs. novel context) or expertise in a given task (Rizzolatti and Sinigaglia 2010) and then related to general and specific predictive aspects. A few studies in humans have, in the past, highlighted the impact of expertise on prediction of movements in groups of experts, such as sport athletes (Aglioti, Cesari et al. 2008), showing their improved ability to anticipate the outcome of a specific action in others. In comparison to sport athletes, musicians possess expertise in both specific sensorimotor skills *and* social signal analyses and exhibit structural brain changes, especially in the temporal and premotor cortex, as a strong biological basis for their expertise and training (Münte, Altenmüller et al. 2002). Abovementioned studies did not specifically test the benefits of expertise onto the recognition of expressive intent in sub-optimal conditions (when information is lacking or altered), although it should be thoroughly studied. In fact, such line of research should critically contribute to the understanding of the behavioral and neural mechanisms underlying a better coordination between several individuals. In sub-optimal conditions, the cerebellum could be highly relied upon in order to predict accurately intention based on action understanding, planning, timing especially due to its fine-tuned connections with the basal ganglia and the cerebral cortex (Caligiore, Pezzulo et al. 2017). In fact, the cerebellum is part of the action observation (Sokolov, Gharabaghi et al. 2010) and voluntary movement (Hülsmann, Erb et al. 2003) networks and it is also a strong hub for timing and rhythm processing (Molinari, Leggio et al. 2007, Chen, Penhune et al. 2008). Cerebellar activity is also enhanced during sensorimotor coordination in musicians (Krause, Schnitzler et al. 2010) and widely recruited by abstract social cognition, with involved sub-regions overlapping with sensorimotor cerebellar territories (Van Overwalle, Baetens et al. 2014, Sokolov 2018). The probability of the involvement of the cerebellum in sub-optimal action and intention decoding is therefore very high.

To shed light on the mechanisms of communicative intent, we recruited musicians and matched control participants who evaluated the visual dynamics of short pieces of violin solos with the violinist represented as a point-light display (PLD) following motion capture recordings on an independent group of expert violinists. Communicative intent was materialized by categorizing Piano vs. Forte intentional gestures in these short musical pieces while undergoing continuous brain scans using functional magnetic resonance imaging (fMRI). These short PLD video were manipulated to include both original (unmodified, but only with visual information) and modified PLD segments: these modified segments included 1) a condition with spatially randomized initial positions of points (namely, the “Spatial shuffling” condition) and 2) a condition in which pieces were cropped to keep only the first instants of the segments (namely, the “Temporal cropping” condition). We then compared the performance of musicians and controls over the accurate recognition of the expressive play on both normal and altered versions of the segments during fMRI. We hypothesized a clear and consistent performance advantage of expert musicians over control participants in assessing communicative intent (Piano vs. Forte). Regarding neuroimaging data, we hypothesized enhanced activity in the frontoparietal network and the cerebellum as a function of expertise that could be modulated by interindividual differences. Finally, we predicted a stronger coupling between the prefrontal and premotor cortex as well as between the cerebellum and the frontoparietal network for experts, i.e. musicians.

## Material and Methods

### Participants

Thirty-seven right-handed participants took part in this study but final sample size was 35 (25 females, 19 experts, 17 female musicians, M age = 27 ± 7 years). In fact, two participants fell asleep repeatedly during data acquisition and were therefore excluded from the final sample. Participants were composed of a group of expert violinists who received at least an 8-yearlong training at the high musical institution (Geneva School of Music) and a control group of non-musicians. While the number of males and females differed significantly between the two groups (χ²(1, N=35) = 4.83, p = 0.027) with more woman violinists than male, age did not differ between the two groups (t(30.26, N = 35) = −0.38, p = 0.7). However, the variance induced by the sex did not seem to impact our models (Supplementary Table 1 & 2). All participants were naive to the experimental procedure and material and had no known history of psychiatric or neurological incidents. Finally, all participants reported normal hearing abilities and normal or corrected-to-normal vision (contact lenses or MRI-compatible plastic glasses). This study was conducted in compliance with the protocol, the current version of the Declaration of Helsinki, the ICH-GCP or ISO EN 14155 (as far as applicable) as well as all legal and regulatory requirements of the State-wise Ethics Committee of the University Hospital of Geneva.

### Experimental Stimuli: Communicative intent task

The complete set of stimuli consisted of 64 videos. These stimuli were produced using the following procedure. First, we filmed using a motion capture system (Qualisys, time sampling) the first violinist of a professional string quartet during 16 rehearsals of the same musical piece, the Death and the Maiden by Schubert, chosen for it offers a wide variety of writing and expressive styles. For half of the rehearsals, the first violinist played alone, for the remaining half, he played with the other string quartet members. During filming, the musicians were instructed to play expressively as if they were giving a concert performance. The recording sessions took place over two days in a concert hall for it offers naturalistic conditions, which perfectly adapt to the musician’s needs and expectations (e.g., quality of the acoustics). After filming, all clips were edited, and the motion capture data were preprocessed to eliminate spurious data through standard filtering process (despiking and smoothing using MATLAB (The Mathworks, Inc., Natick, MA, USA)) in order to produce cleaned point-light displays (PLD) of the performances. The next step was to select specific moments in each performance where the first violinist indicates his intent to the fellow musicians with two communicative intents: piano and forte. To do so, we defined, together with the musicians, these key moments in the music score and edited the corresponding 16 short sequences (8 Piano, 8 Forte). Two experimental visual manipulations were then applied on each edited point-light sequence. The first manipulation consisted in segmenting the sequence in two parts: the first part referred to the preparation of the entry, just before the production of the Piano or Forte gesture;1.2*s*), the second one referred to the whole sequence (i.e., movement preparation plus entry;2.7*s*). We refer to these sequences as Temporally cropped vs. Temporally unmodified. In the temporally cropped sequences, we focused on attack sequence because musicians think that it is crucial to understand the musicians’ specific capacity to coordinate with one another. The second manipulation consisted in destroying the anthropomorphic shape of the stimuli through a single spatial scrambling process, which keeps the dynamics of each point but shuffles their relative relationships with others. The final shape, which keeps the same kinetic energy as the original, does not related to any anthropomorphic shape. We refer to these sequences as Spatially shuffled vs. Spatially unmodified. They were designed to highlight musicians’ higher processing of dynamic visual information when visual bodily anthropomorphic references are missing.

The final excerpts presented to the participants were a combination of all three conditions in a pseudorandom order: the communicative intent, the temporal cropping, and the spatial shuffling. For example, an excerpt could be presenting a piano gesture, temporally cropped, but spatially unmodified while another one could be presenting a forte gesture, with temporal cropping and spatial shuffling. In all sequences, only visual information were used and the accompanying sound tracks were not presented (see an example of each condition in Fig.1).

**Fig.1:**
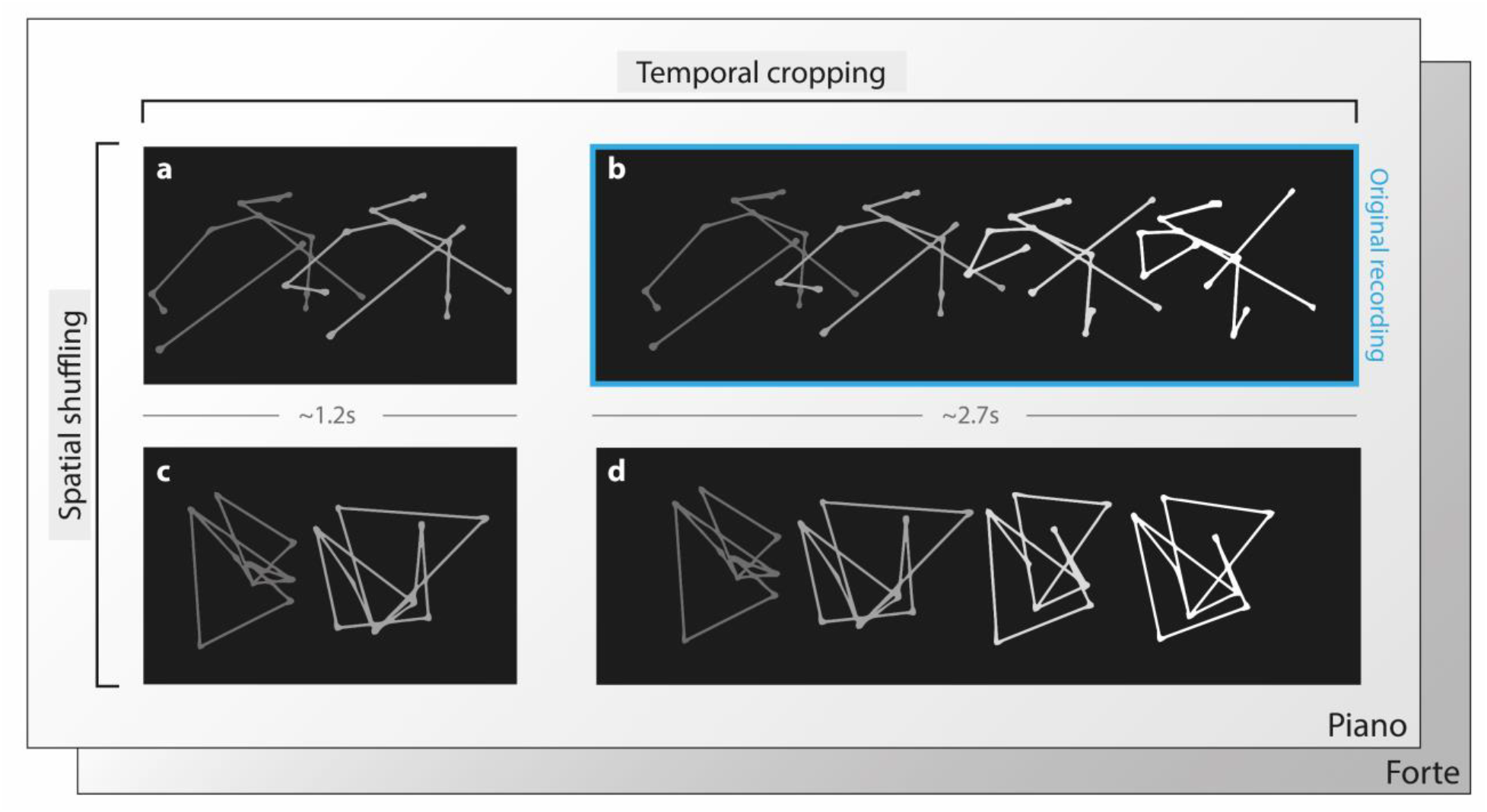
Overview of the stimuli displayed to the participant. The sequence of expressive movement is displayed through point-light display based on the collected motion capture of a string quartet first violinist. The two temporal segments used in the experiment respectively refer to the preparation (Temporally cropped condition, a) and the preparation plus the entry part (Temporally unmodified condition, b). For each of these temporal conditions, spatial shuffling was also applied as a separate condition (c, d, respectively). After watching the selected sequence, the participants were asked to indicate the perceived communicative intent of the music (forte or piano) by pressing a button.

### Experimental procedure: Communicative intent task

Participants, divided in two groups (musician vs control participants) were subjected to a 2 (communicative intents: Piano vs. Forte) × 2 (temporal cropping: Temporally cropped vs. Temporally unmodified) × 2 (spatial shuffling: Spatially shuffled vs. Spatially unmodified) within-subject factorial design, resulting in eight experimental conditions (Fig.1). Before scanning, participants were introduced to and familiarized to the experimental task. This included performing two trials from each condition (these stimuli were excluded from the MRI task) while the experimenter monitored performance. During the MRI session, participants watched the point-light display movies of the violinist movements. After each trial, participants had to report whether they thought whether the performance is related to a communicative intents: forte or piano. During debriefing, all participants reported complete comprehension of the experimental task. The experimental procedure relied on a two-alternative-forced-choice task. It aimed at revealing the difference in optimal decision strategies (Gold and Shadlen 2002, Ratcliff and McKoon 2008, Deneve 2012) developed by groups of musicians (task experts) versus control participants (non-experts).

### Behavioral Data Analysis

The statistical software R was used to analyze all behavioral data. We computed generalized linear mixed model (GLMM) to estimate the variance explained by the fixed factors Piano/Forte and Musician/Controls on the percentage of correct responses. GLMM takes advantage of the modelling of random effects to improve precision of the model and allow for the computation of models with non-normal distribution, here a binomial distribution. We test our predictions for the effect of different fixed effect factors including the expertise of the participant, the communicative intents, the spatial shuffling, and the temporal cropping. The random intercept effects encapsulated the variability related to each participant. We used a step-up strategy while building the model to test the different combinations of fixed effects. Based on the marginality principle, we present the highest order interaction effects (Nelder 1977), namely the interaction between the expertise and the other aforementioned experimental conditions. We used chi-square difference tests to investigate the contribution of each variable and their interaction. We report the effect sizes in accordance with the approach of Nakagawa and Schielzeth, implemented in the “MuMIn” R package (Nakagawa and Schielzeth 2013). They created an approach based on two indicators, a marginal and a conditional R2 (respectively, R2m and R2c). R2m is the variance explained by the fixed factors, whereas R2c is the variance explained by the entire model (both fixed and random effects). These two indicators allows comparability with standard methods, while taking into account the variance explained by the random effects. We calculated and reported them for each statistical models.

### Neuro-imaging Image Acquisition

Imaging data were acquired using a Siemens Trio 3.0 tesla MRI scanner at the Brain and Behavioral Laboratory (BBL) of the University Medical Center, University of Geneva (Geneva, Switzerland). For each participant and for the run of the experimental task we acquired 290 functional T2* -weighted echo planar image volumes (EPIs; slice thickness=3mm, gap=1mm, 36 slices, TR=650ms, TE=30ms, flip angle = 90°, matrix=64·64, FOV=200mm). A T1-weighted, magnetization-prepared, rapid-acquisition, gradient echo anatomical scan (slice thickness=1mm, 176 slices, TR=2530ms, TE=3.31ms, flip angle = 7°, matrix=256·256, FOV=256mm) was also acquired. Therefore, for each participant we acquired 290 volumes of 36 slices for a total of 10’440 slices. The grand total for all participants was 10’150 volumes and 365’400 slices.

### Neuro-imaging Data Analysis

Functional data were analyzed using Statistical Parametric Mapping version 12 (https://www.fil.ion.ucl.ac.uk/spm) operating in MATLAB (The Mathworks, Inc., Natick, MA, USA). Preprocessing steps included realignment to the first volume of the time-series to correct for head motion; slice-timing; normalization to the Montreal Neurological Institute (MNI) template (resampled at 3 × 3 × 3 mm) and eventually spatial smoothing with an isotropic Gaussian kernel of 8 mm full width at half-maximum. A high-pass filter of 128s was used to remove low frequency components. Then, a general linear model (first-level analysis) was defined for each participant separately (within-subject statistics). For the experimental task, correctly-evaluated trials were modeled by specific boxcar functions defined by the duration of the video stimuli spanning stimulus onset to offset and convolved with the canonical hemodynamic response function. Group-level statistics were then performed using a flexible factorial design to take into account the variance of all conditions and participants. Two different group-level models were computed for the present data: Model 1 included eight conditions (1. Spatially unmodified, Temporally unmodified, and Piano; 2. Spatially unmodified, Temporally unmodified, and Forte; 3. Spatially unmodified, Temporally cropped, Piano; 4. Spatially unmodified, Temporally cropped, Forte;5. Spatially shuffled, Temporally unmodified, Piano; 6. Spatially shuffled, Temporally unmodified, and Forte; 7. Spatially shuffled, Temporally cropped, and Piano; 8. Spatially shuffled, Temporally cropped, and Forte) and two groups (Musician; Control) with no covariates [Group × Conditions] while Model 2 included task performance as a group-level covariate of interest interacting with the Conditions factor [Performance × Conditions]. Both models additionally included a mandatory “participant factor” that allows inter-subject variability calculation. For both group-level models, specification of independence was set to true for the Participant and Group factors while it was set to false for the other factor conditions. Regarding variance estimation, it was set to unequal for all factors including Group, as homoscedasticity criteria can usually not be met with fMRI data (default setting in SPM12). For both models, group-level results were voxel-wise thresholded in SPM12 using corrected statistics with *p*<.05 False Discovery Rate (FDR) and an arbitrary cluster extent threshold of k>10 voxels. For all analyses, regions with significant activation enhancement were labeled based on probabilistic cytoarchitechtonic atlases (Automated Anatomical Labelling atlas (Tzourio-Mazoyer, Landeau et al. 2002), Cerebellum atlas (Diedrichsen, Balsters et al. 2009, Diedrichsen, Maderwald et al. 2011)) and rendered on semi-inflated brains of the CONN toolbox (http://www.nitrc.org/projects/conn), see Fig.2.

**Fig.2:**
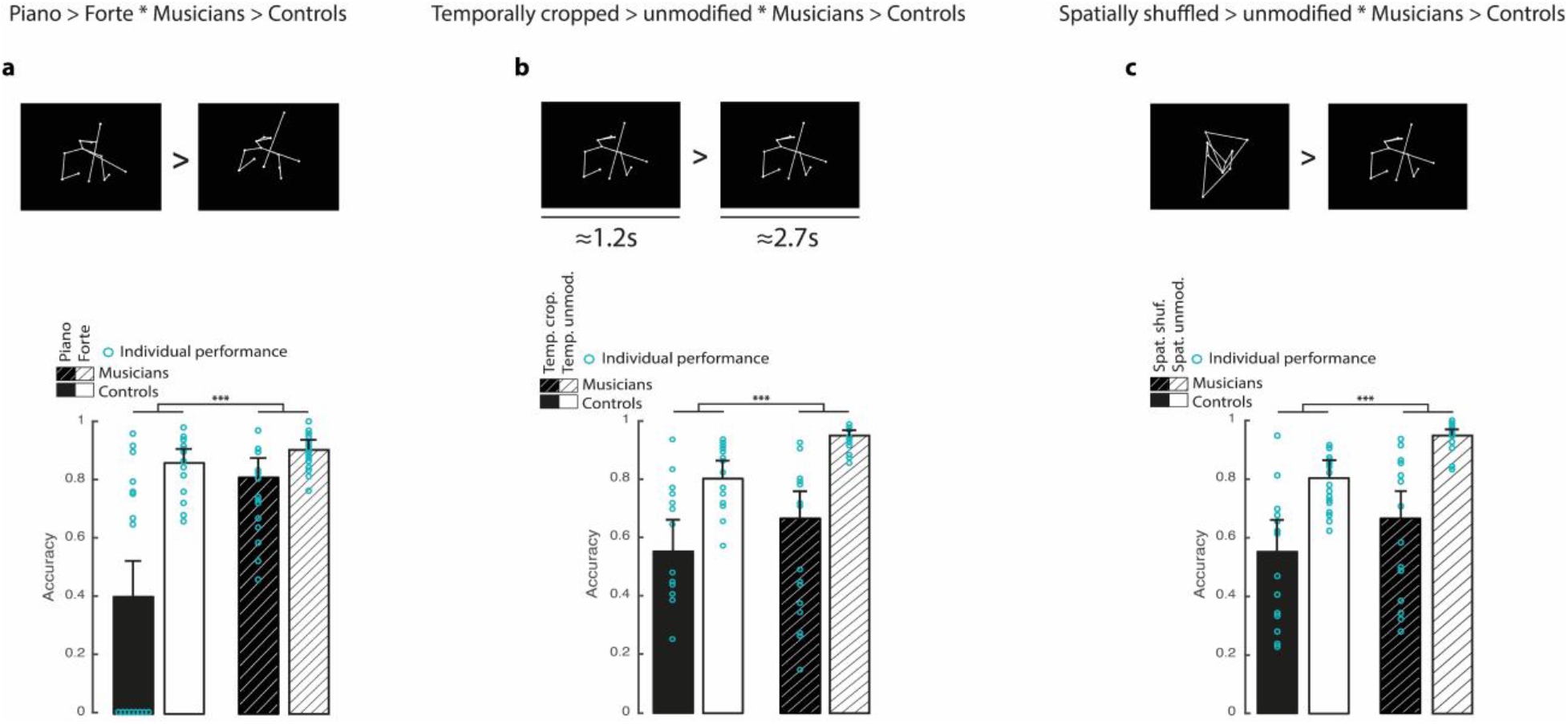
Experimental stimuli and behavioral results for the impact of expertise on intention evaluation. (**a**) Example of piano vs. forte Point Light Display (PLD) and averaged performance and individual points per group for piano vs. forte piece dynamics. (**b**) Example of temporally cropped vs. temporally unmodified PLD and averaged performance and individual points per group for piano vs. forte piece dynamics in temporally cropped vs. temporally unmodified sequences. (**c**) Example of spatially shuffled vs. spatially unmodified PLD and averaged performance and individual points per group for piano vs. forte piece dynamics in spatially shuffled vs. spatially unmodified sequences. Error bars represent 95% confidence interval. [***: *p*<.001, **: p<.01, *: p<.05; : p<0.1; Spat.: spatial, Temp.: temporal, shuf.: shuffling, crop.: cropping, unmodif.: unmodified]

#### Model 1: Conditions Training group-level statistics

For this first model, regressors were created for each experimental condition and for each participant (N=35), leading to a first-level design matrix including 16 regressors in total among which 8 regressors of interest (Conditions, see above) and 8 regressors of no interest (Incorrect trials, the six motion parameters and the constant term). Each regressor of interest was used to compute main effect contrast vectors that were then taken to a second-level, group analysis using the flexible factorial design specification that we will detail here. Group-level analysis included the following factors: Conditions (see above) and Group. The Conditions factor was used to compare the musicians to the control participants regarding their ability to assess the communicative intent of the stimuli, whether spatially shuffled and/or temporally cropped. The following contrasts were hence computed using the above factorial architecture of the data: [piano > forte * musicians > controls], [temporally cropped > temporally unmodified *Musician>Control] and [spatially shuffled>spatially unmodified*Musician>Control] (see Fig.2).

#### Model 2: Conditions Training group-level statistics with task performance as second-level covariate

The second model used the exact same settings and factorial structure as Model 1 in addition to a group-level covariate taking into account task performance for each participant (percentage of hits during the experimental task). This covariate was set to interact with the Conditions factor. Therefore, Model 2 included the following factorial structure: Conditions*Task performance. This model was used to constrain our statistical results and observe the way some brain regions may be sensitive to task performance in Musicians as opposed to Non-musicians participants for the following contrasts : [piano > forte * performance], [temporally cropped > temporally unmodified * performance] and [spatially shuffled > spatially unmodified * performance]. Results from this second model were overlaid in green-to-blue in Fig.2.

### Functional connectivity analyses

Functional connectivity analyses were carried out using the CONN toolbox v18.a (Whitfield-Gabrieli and Nieto-Castanon 2012). Spurious sources of noise were estimated and removed using the automated toolbox preprocessing algorithm, and the residual BOLD time-series was band-pass filtered using a low frequency window (0.008 < f < 0.09 Hz). Correlation maps were then created for each condition of interest by taking the residual BOLD time-course for each condition from atlas ROIs and computing bivariate Pearson’s correlation coefficients between the time courses of each voxel of each region of the atlas. These correlations were then converted to normally distributed values using Fisher’s transform. Finally, group-level analyses were performed on these Fisher-transformed correlation maps to assess for main effects within-groups and significant connectivity differences between groups for contrasts of interest. Type I error was controlled by the use of seed-level false discovery rate (FDR) correction with *p*<.05 to correct for multiple comparisons.

## Results

### Behavior

Behavioral results highlighted a generalized and reliable advantage of musicians over control participants for accurately evaluating piece dynamics as expressing either Piano or Forte communicative intentions in a discriminative task (Fig.2). We tested how the interaction effects of our factors (Group [Musicians>Controls] and Conditions [Forte>Piano, Temporally cropped>Temporally unmodified, Spatially shuffled>Spatially unmodified]) explained a larger part of variance for each statistical model compared to models with only the main effects (all *p*<.001, complete statistics in Supplementary Table 2). More specifically, for each model computed, we observed that when information is altered (Temporally cropped, Spatially shuffled; Fig.2**b,c**) or subtler (Piano, Fig.2**a**), the performance of all our participants drops significantly (all *p*<.001, complete statistics in Supplementary Table 2 & 3). At the group level, musicians outperformed control participants in assessing communicative intents regardless of the condition presented (Fig.2**a-c**, all *p*<.001). Finally, we describe a significant interaction effect between group and conditions for each model (all *p*<.001). No sex differences were observed (Supplementary Figure 1, Supplementary Table 1).

### Neuroimaging

#### Wholebrain data

Neuroimaging results focused on the regions with enhanced activations for musicians as compared to control participants, for communicative intent, temporal cropping and spatial shuffling conditions compared to complete excerpts. Analyses focused exclusively on trials in which participants correctly identified the intention depicted in the PLD. When focusing on the excerpts expressing a piano nuance (Piano>Forte*Musician>Control), we observed enhanced activations in the pre supplementary area (preSMA) (MNI coordinates in Supplementary Table 4 & 5) and left dorsolateral prefrontal cortex (DLPFC) (Fig.3**a,j**, Supplementary Table 4 & 5, Supplementary Figure 2 & 3). When focusing on the temporally cropped excerpts (Temporally cropped>Temporally unmodified*Musician>Control), we observed enhanced activations in the left IPL, cerebellum lobules V, VI, VIIb, and VIII, the bilateral posterior middle temporal gyrus (MTG), and the inferior frontal gyrus (IFG) (Fig.3**b,c,f,g,k,l**; Supplementary Table 4 & 5, Supplementary Figure 2 & 3). Finally, when we focused on the spatially shuffled excerpts (Spatially shuffled>Spatially unmodified*Musician>Control), we observed enhanced activations in the bilateral intraparietal sulcus (IPS), right preSMA, and bilateral DLPFC extending to the IFG pars opercularis and triangularis, cerebellum (Vermis areas IV,V, Crus II, lobules I-IV, V, VI, VIIb, VIIIa, VIIIb, X), and bilateral insula (Fig.3**d,e,h,i,m,n**; Supplementary Table 4 & 5, Supplementary Figure 2 & 3). To further emphasize these enhanced activations in perspective with the global individual performance of each participant, we also computed another second-level analysis with individual performances of the participants as group-level covariate. Consequently, we could account for brain regions with enhanced activations in relation to general performance (group-level analysis) for our contrasts of interest. These results crucially include inter-individual variability regarding task performance, across groups (continuous, one average value per participant). Rendered in green-to-blue activations (Fig.3**b-e,f-i,k-n**; Supplementary Table 4 & 5, Supplementary Figure 2), global individual performance analysis revealed a strong overlap with abovementioned brain regions for the temporally cropped condition, especially in the IPL, SMA, and in the cerebellum (Temporally cropped>Temporally unmodified*Musician>Control with Individual Performance, Fig.3**b,c,f,g,k,l**; Supplementary Table 4 & 5, Supplementary Figure 2). This result suggests that this network is highly relied on for accurately evaluating the communicative intent of temporally cropped PLD as a function of both performance and expertise and is sensitive to individual differences of performance as well. Such overlap between analyses was smaller for the spatially shuffled excerpts (Spatially shuffled>Spatially unmodified*Musician>Control with Individual Performance), especially in the cerebellum. In fact, results show that portions of the bilateral insula, bilateral preSMA, bilateral DLPFC as well as the right IPS did overlap, emphasizing the strong role of these regions in both performance and expertise. Interestingly, the left IPS showed a much smaller overlap between analyses compared to the right IPS, suggesting an interhemispheric dissociation (Fig.3**d,e,h,i,m,n**) with left IPS activity enhanced massively for correctly evaluated piece dynamics as opposed to the right IPS being modulated by the interindividual differences of performance. Such overlap between analyses was also observed bilaterally for cerebellum area VIIb (Fig.3**m**). Additional brainstem activity was observed in the right corticospinal tract (Fig.3**n**).

**Fig.3:**
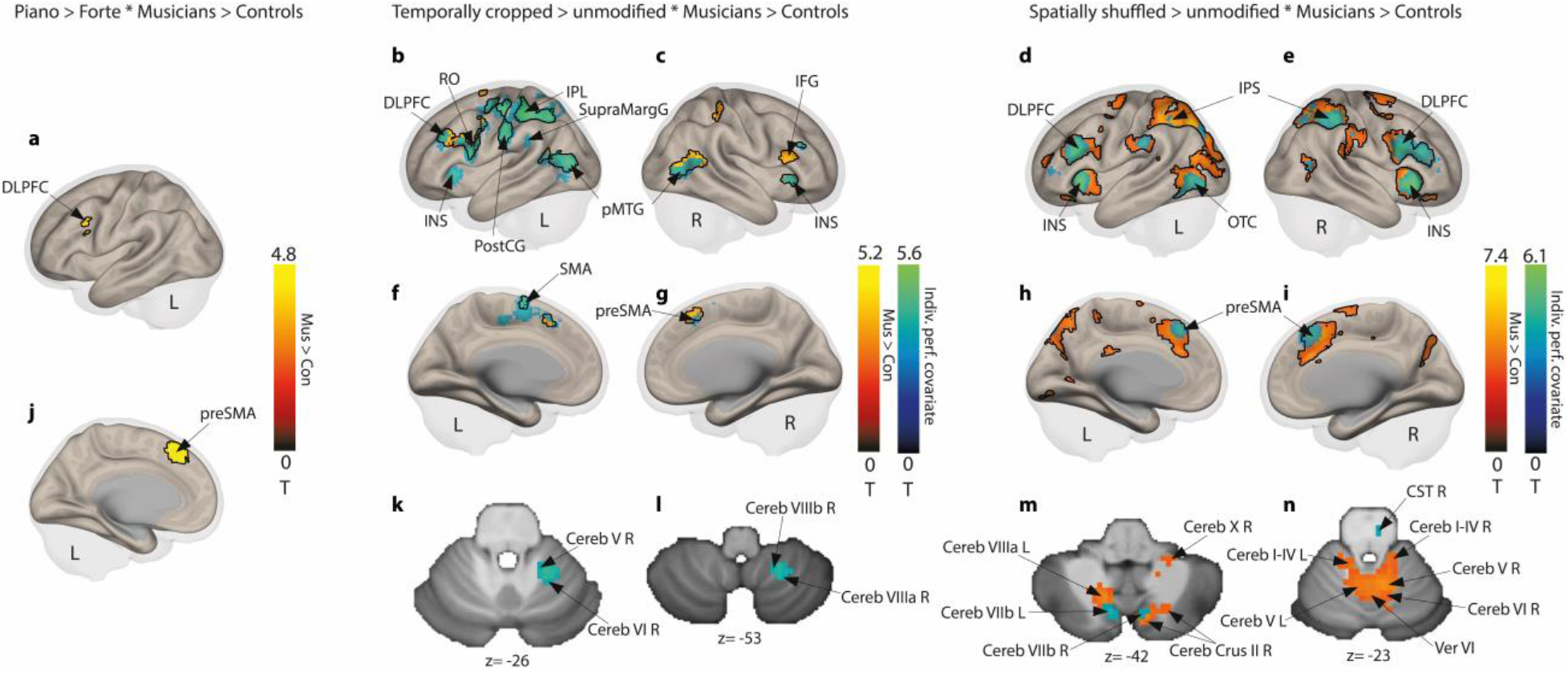
Neural evidence of communicative intent decoding by musicians and matched controls. (**a,j**) Enhanced activity in preSMA and DLPFC for piano vs. forte sequences for musicians vs. control participants. (**b,c,f,g**) Enhanced activity for temporally cropped vs. temporally unmodified sequences for musicians vs. control participants in red-to-yellow in IPL, and with overall task performance as group-level covariate in blue-to-green in IPL, DLPFC, pMTG, preMSA. and in subregions of the cerebellum (**k,l**). (**d,e,h,i**) Enhanced activity for spatially shuffled vs. spatially unmodified sequences for musicians vs. control participants in red-to-yellow in IPS, preSMA, DLPFC, and INS, and with overall task performance as group-level covariate in blue-to-green in IPS, preSMA, DLPFC, INS, and OTC and in the cerebellum (**m,n**). The color bars represent the statistical T values of the contrast. Black outlines delineate regions of the contrasts without the performance covariate. [Cereb: cerebellum lobule; Cereb Crus: cerebellum crus of ansiform lobule; CST: corticospinal tract of the brainstem; DLPFC: dorso lateral prefrontal cortex; IFG: inferior frontal gyrus; INS: insula; IPL: inferior parietal lobule; IPS: inferior parietal sulcus; lingual gyrus; OTC: occipito-temporal cortex; pMTG: medial temporal gyrus; posterior part; preSMA: pre supplementary motor area; PostCG: postcentral gyrus; RO: Rolandic operculum; SupraMargG: supra marginal gyrus; Ver: vermis; L: left; R:right]. Voxel-wise p<.05 FDR corrected.

#### Functional connectivity data

Additionally, atlas-based seed-to-seed functional connectivity (FC) analyses were performed to highlight the existence of widespread coupled brain activity targeting frontoparietal and cerebellar regions and related to both communicative intent processing and expertise. These analyses revealed the involvement of numerous regions observed in our wholebrain contrasts of interest in addition to subcortical and cerebellar connectivity (Fig.4). A general effect of expertise across conditions (Musician>Control, main effect of all conditions) revealed a functional coupling between the left IPL and the left post central gyrus (Supplementary Figure 4). Concerning our contrasts of interest, communicative intent and expertise interacted (Piano>Forte*Musician>Control) and lead to both coupled and anti-coupled functional networks (Fig.4**a**; Supplementary Table 6). Specifically, we observed coupled FC between the bilateral MTG, left putamen, bilateral fusiform cortex, brainstem, and several subregions within the cerebellum, such as cerebellar lobules III, VIII and × of the left hemisphere. Anti-coupled FC was observed between the medial frontal cortex, posterior cingulate gyrus, frontal pole, left DLPFC, and left IPS (see details in Fig.4**a**; Supplementary Table 6). For temporally cropped excerpts (Temporally cropped>Temporally unmodified*Musician>Control), only coupled FC was observed. More specifically, analyses showed widespread fronto-parieto-cerebellar FC in the bilateral inferior frontal gyrus pars opercularis (IFGop), left DLPFC, left superior parietal lobule (SPL), right IPS and vermis areas VII and VIII (see details in Fig.4**b**; Supplementary Table 6). The last contrast of interest involving visually shuffled PLD (Spatially shuffled>Spatially unmodified*Musician>Control) highlighted coupled FC between the anterior part of the left inferior, middle and superior temporal gyri (aITG, aMTG and aSTG, respectively), left posterior MTG and right supramarginal gyrus while anti-coupled FC characterized connectivity between the left IFGop, right posterior ITG, posterior cingulate gyrus and brainstem (see details in Fig.4**c**; Supplementary Table 6).

**Fig.4:**
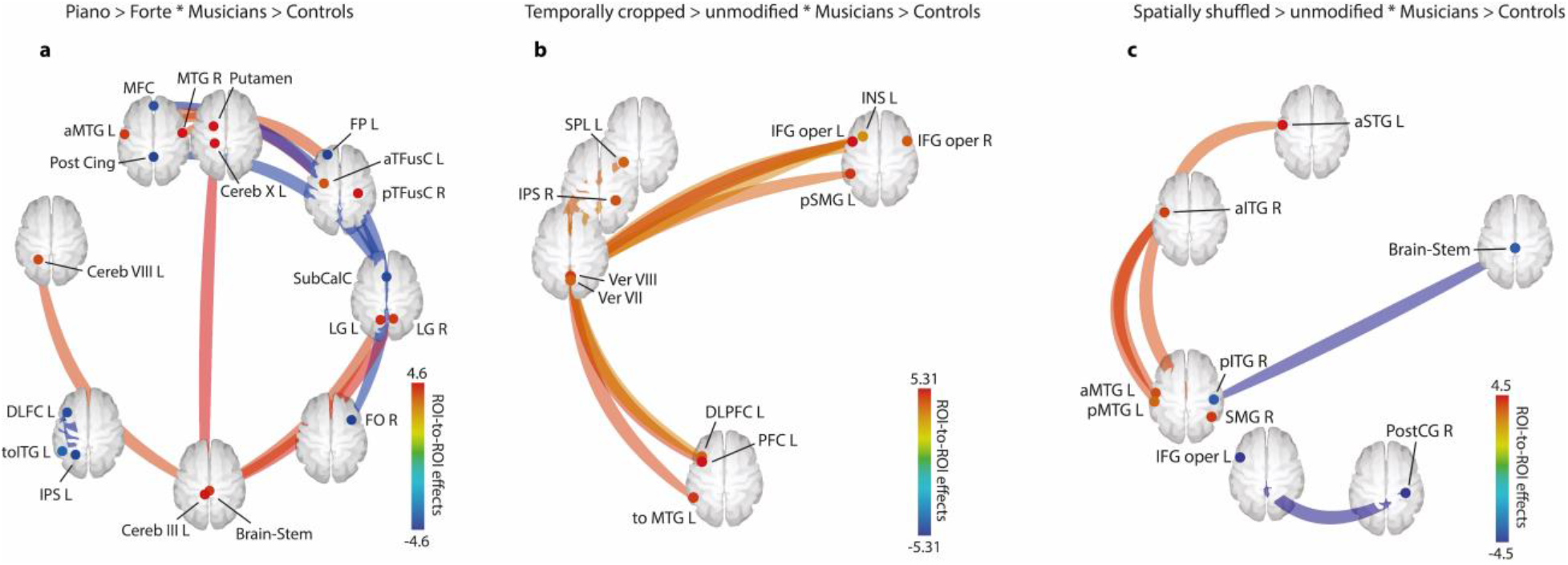
Functional connectivity of communicative intent decoding by musicians and matched controls. (**a**) Enhanced connectivity for piano vs. forte excerpts for musicians vs. control participants. (**b**) Enhanced connectivity for temporally cropped vs. unmodified sequences for musicians vs. control participants. (**c**) Enhanced connectivity for spatially shuffled vs. unmodified sequences for musicians vs. control participants [aITG: inferior temporal gyrus, anterior part; aMTG: medial temporal gyrus, anterior part; aSTG: superior temporal gyrus, anterior part; aTFus: temporal fusiform, anterior part; Cereb: cerebellum lobule; DLPFC: dorso lateral prefrontal cortex; FO: frontal operculum; FP: frontal pole; IFG oper: inferior frontal gyrus operculum; INS: insula; IPL: inferior parietal lobule; IPS: inferior parietal sulcus; LG: lingual gyrus; MFC: medial frontal cortex; MTG: medial temporal gyrus; PFC: prefrontal cortex; pITG: inferior temporal gyrus, posterior part; pMTG: medial temporal gyrus, posterior part; Post Cing: posterior cingulate; PostCG: posterior central gyrus; pSMG: superior medial gyrus, posterior part; pTFusC: temporal fusiform cortex, posterior part; R: right; SMG: superior medial gyrus; SPL: superior parietal lobule; SubCalC: subcallosal cortex; toITG: inferior temporal gyrus temporo-occipital part; toMTG: medial temporal gyrus, temporo-occipital part; Ver: vermis]. Seed-level p<.05 FDR corrected.

## Discussion

The present study aimed at a clearer understanding of the interaction between expertise and communicative intent evaluation as a potential proxy for social interactions and coordination. Using point-light displays of violinists as stimuli, we asked experts and non-experts, namely musicians and controls, to evaluate the expressive intent of the plays. Expressive intent could materialize as *piano* or *forte* and either be visually or temporally unmodified or shuffled and cropped, respectively. Our results demonstrate that musicians consistently outperformed control participants, be it for unmodified or altered stimuli. Pre-motor and lateral parietal areas together with dorsolateral prefrontal cortex played a major role in giving a strong hand to experts in addition to numerous cerebellar regions.

Behavioral results confirmed and further developed the role of expertise in the perception of others’ intent. Musicians were more consistent and accurate in perceiving expressive gestures, indicating close links between perception and action abilities (Wöllner and Cañal-Bruland 2010). Importantly, such result remained true when information was lacking or altered, highlighting behavioral benefits of expertise in action understanding, integration, and prediction based on the short dynamics of the segments and even with altered anthropomorphic information. It stresses the importance for action processing and prediction of combining perceptual information about the observer’s own and the others’ goal-directed movements (Spunt and Lieberman 2012).

Our neuroimaging results revealed a widespread network of brain regions as a function of expertise when decoding or inferring communicative intent, especially in strongly altered perceptual conditions. The advantage of experts for understanding and assessing communicative intent recruited the DLPFC, associated with observation of actions (Rizzolatti and Sinigaglia 2010) and influenced by training and expertise (Moore, Cohen et al. 2006) as well as the preSMA, involved most notably in internally and externally selected actions (Mueller, Brass et al. 2007). Therefore, preSMA and DLPFC seem sufficient to accurately internalize action and extract communicative intent, respectively, in expert participants. These regions would also explain the ability of musicians to integrate the temporal structure of rhythm mediated by working memory (Chen, Penhune et al. 2008). Functional connectivity analyses also revealed that expert participants relied on both coupled and anti-coupled networks to successfully infer communicative intent. The coupled networks include areas involved in intention probability (putamen; Zapparoli, Seghezzi et al. 2018) as well as in movement prediction and motor imagery (cerebellum; Sokolov, Miall et al. 2017) and timing encoding (brainstem and cerebellum; Rao, Mayer et al. 2001, Molinari, Leggio et al. 2007). More specifically, a large region of the cerebellum including lobule VIII (and VIIIa) was shown to covary with instrumental expertise, especially for temporal complexity (Chen, Penhune et al. 2008). This result raises the question of whether general or instrument-specific abilities of musicians directly influence cerebellum activity, such important distinction should be addressed in future studies. In a lesion study, lobule III—as well as lobules I, IV and V—was also involved in action observation (Sokolov, Gharabaghi et al. 2010) while lobule IX and to a smaller extent lobule × were repeatedly linked to verbal working memory (Van Overwalle, Baetens et al. 2014). As a functionally connected cerebellar network and through connections with the brainstem and basal ganglia, these lobules therefore give strong weight to the cerebellum as a crucial contributing player in action observation, processing, and prediction. On the other hand, anti-coupled activity recruited brain regions known for mental states attributed to moving shapes and memory for intentions (Medial prefrontal cortex and frontal pole, respectively; Blakemore and Decety 2001), action observation (DLPFC: Rizzolatti and Sinigaglia 2010), and biological motion perception (ITG; Blakemore and Decety 2001), as well as attention to intention and its understanding (IPS; Blakemore and Decety 2001).

Behavioral advantage of communicative intent assessment in altered perceptual conditions by expert participants relied essentially on several of the abovementioned brain areas with some additions. In temporally cropped sequences, expertise further recruited positively connected regions involved in interoception and motor intention awareness (insula; Craig and Craig 2009), intentional action production (IFGop; Zapparoli, Seghezzi et al. 2018), intention understanding (IPS and IPL; Blakemore and Decety 2001) and temporal processing linked to actions (vermis, especially areas VIII and VIIIa; Rao, Mayer et al. 2001). Communicative intent understanding as a function of expertise in spatially shuffled PLD recruited very similar brain areas when compared to temporal cropping at the wholebrain level, but with larger clusters. This result could be explained by an advantage in assessing complex visual inputs in musicians as compared to non-musicians to evaluate instrumental performance (Griffiths and Reay 2018). Moreover, general task performance as group-level covariate constrained our wholebrain results and showed a difference between left and right IPS, with the latter showing a larger cluster of enhanced activity as a function of task performance in experts vs. controls. This suggests a specific role of the right IPS region related to inter-individual differences in performances and this region was indeed reported to contribute to interpersonal synchronization related to action (Bhat, Hoffman et al. 2017). Additionally, functional connectivity analyses revealed coupled temporal cortices (anterior STG, MTG, ITG; posterior MTG, ITG) and anti-coupled connectivity in the IFGop, brainstem and posterior cingulate cortex. The coupled networks revealed processing of biological motion independent of motor information (superior and middle temporal cortices; Rizzolatti and Sinigaglia 2010) and attribution of intentions to spatially shuffled stimuli (posterior STG; Lee, Gao et al. 2012), regions interestingly known for their feedforward connections to the IPS and IPL (Rizzolatti and Sinigaglia 2010). Anti-coupled functional connectivity stressed again intentional action production and respectively unusual action intention processing (IFGop; Zapparoli, Seghezzi et al. 2018) and movement initiation and control (brainstem (Nandi, Aziz et al. 2002) and cerebellum (Rao, Mayer et al. 2001)).

While our data shed new light on intention encoding and decoding and further highlight behavioral contexts favorable to expert participants over non-experts, some limitations should be taken into account. First, sample size could have been larger, although it was difficult to recruit several additional musicians who would fit our inclusion criteria. Second, past studies emphasized structural brain differences between musicians and non-musicians (Gaser and Schlaug 2003) and therefore we could constrain our results even more had we acquired diffusion tensor images, for instance. Third, our stimuli included point-light displays of violinists only, therefore restricting our conclusions regarding other types of expert musicians. Finally, other ways of altering the stimuli, such as modifying rhythmicity or adding sublevels of visual shuffling, could have been employed to further specify the impact of expertise on more fine-grained perceptual alterations and their impact on communicative intent decoding.

Taking our behavioral and neuroimaging data as well as study limitations into consideration, our results reveal the strong role of expertise in understanding and predicting action and communicative intentions. Such assertion is especially true in altered perceptual conditions, namely with visually or temporally altering of stimuli, and this advantage of musicians over non-musicians would rely on regions of the frontoparietal network in addition to several areas of the cerebellum, basal ganglia and brainstem. Such neural systems might be at play in numerous other conditions in everyday social interactions as humans are experts in predicting others when they interact with them daily.

## Acknowledgement

This study was supported by a Swiss National Science Foundation project funding (51NF40-104897 – DG). The EU ICT SIEMPRE project acknowledges the financial support of the Future and Emerging Technologies (FET) programme within the Seventh Framework Programme for Research of the European Commision, under FET-Open grant number 250026-2. We thank Bruno Bonet at the Brain and Behavior Laboratory of the Geneva University Hospital Medical Center for his assistance during data acquisition. Finally, we would like to extend our acknowledgements to musicians from the Quartetto di Cremona for the recording of the stimuli, and the contribution of violinist experts Chiara Noera and Florence Malgoire for their precious suggestions.

## Data availability statement

The datasets generated during and/or analyzed during the current study are available from the corresponding author on request.

## Code availability statement

The codes used to analyze the data of the current study are available from the corresponding author on request.

## Notes

### Competing Interest Statement

The authors have declared no competing interest.

