## Supplementary material for "Frontoparietal, cerebellum network codes for accurate intention prediction in altered perceptual conditions"

#### Supplementary Table 1

**Model statistics**

| Model (participant and sex as random effect) | Information Criterion | | Statistics | | | |
|---|---|---|---|---|---|---|
| | AIC | BIC | Chisq(1, N = 35) | Pr(FDRadj) | $R^2_m$ | $R^2_c$ |
| Expertise*Nuance | 5916.6 | 5957.4 | 107.76 | <0.001 | 0.18 | 0.38 |
| Expertise+Nuance | 6022.3 | 6056.4 | | | | |
| Expertise*Temporal cropping | 6265.8 | 6306.7 | 84.964 | <0.001 | 0.16 | 0.32 |
| Expertise+Temporal cropping | 6348.7 | 6382.8 | | | | |
| Expertise*Spatial shuffling | 6120.7 | 6161.6 | 60.479 | <0.001 | 0.21 | 0.37 |
| Expertise+Spatial shuffling | 6179.2 | 6213.2 | | | | |

Each model encompassing the interaction effect between the expertise and one of the three conditions outperformed the corresponding model with only the main effect. The random intercept effect captures the variance induced by every participant and every sex. This indicates the importance of the interaction effect in characterizing the variance of the dataset.

### Supplementary Table 2

**Model statistics**

| Model (participant as random effect) | Information Criterion | | Statistics | | | |
|---|---|---|---|---|---|---|
| | AIC | BIC | Chisq(1, N = 35) | Pr(FDRadj) | $R^2_m$ | $R^2_c$ |
| Expertise*Nuance | 5914.6 | 5948.6 | 107.76 | <0.001 | 0.18 | 0.38 |
| Expertise+Nuance | 6020.3 | 6047.6 | | | | |
| Expertise*Temporal cropping | 6263.8 | 6297.8 | 84.964 | <0.001 | 0.16 | 0.32 |
| Expertise+Temporal cropping | 6346.7 | 6374 | | | | |
| Expertise*Spatial shuffling | 6118.7 | 6152.7 | 60.479 | <0.001 | 0.21 | 0.37 |
| Expertise+Spatial shuffling | 6177.2 | 6204.4 | | | | |

Each model encompassing the interaction effect between the expertise and one of the three conditions outperformed the corresponding model with only the main effect. The random intercept effect captures the variance induced by every participant. This indicates the importance of the interaction effect in characterizing the variance of the dataset.

#### Supplementary Table 3

**Contrast statistics**

| Model | Conditions | Statistics | |
|---|---|---|---|
| | | Chisq( 1, N = 35) | Pr(FDRadj) |
| Expertise*Nuance | Piano (Mus. > Cont.) | 39.578 | <0.001 |
| Expertise*Nuance | Forte (Mus. > Cont.) | 4.9037 | 0.027 |
| Expertise*Nuance | Cont. (Piano > Forte) | 369.59 | <0.001 |
| Expertise*Nuance | Mus. (Piano > Forte) | 17.186 | <0.001 |
| Expertise*Nuance | Mus. > Cont. & Piano > Forte | 107.83 | <0.001 |
| Expertise*Temporal cropping | Temp. crop. (Mus. > Cont.) | 10.484 | 0.001 |
| Expertise*Temporal cropping | Temp. unmod. (Mus. > Cont.) | 47.993 | <0.001 |
| Expertise*Temporal cropping | Cont. (Temp. crop. > unmod.) | 7.5669 | 0.006 |
| Expertise*Temporal cropping | Mus. (Temp. crop. > unmod.) | 87.865 | <0.001 |
| Expertise*Temporal cropping | Mus. > Cont. & Temp. crop. > unmod. | 82.462 | <0.001 |
| Expertise*Spatial shuffling | Spat. shuf. (Mus. > Cont.) | 14.29 | <0.001 |
| Expertise*Spatial shuffling | Spat. unmod. (Mus. > Cont.) | 47.935 | <0.001 |
| Expertise*Spatial shuffling | Cont. (Spat. shuf. > unmod.) | 33.347 | <0.001 |
| Expertise*Spatial shuffling | Mus. (Spat. shuf. > unmod.) | 179.04 | <0.001 |
| Expertise*Spatial shuffling | Mus. > Cont. & Spat. shuf. > unmod. | 57.604 | <0.001 |

Contrasts between the different conditions, groups, and their interaction in each computed model. The contrasts and associated figures (Fig 1) show that musicians outperformed control participants in accurately recognizing piano and forte across all conditions. It also highlights the difficulties in recognizing piano nuance, temporally cropped excerpts, and spatially shuffled excerpts.

#### Supplementary Table 4

**Staistics: p-values adjusted for search volume**

| Model | Region of interest (MNI mm [x,y,z]) | Cluster size (k) | peak-level T-test |
|---|---|---|---|
| Expertise*Nuance | preSMA (0, 27, 45) | 125 | t(1,35) = 5.17, p = 0.005 |
| Expertise*Nuance | DLPFC (-42, 12, 30) | 15 | t(1,35) = 4.11, p = 0.019 |
| Expertise*Spatial shuffling | L INS (-27, 27, -3) | 652 | t(1,35) = 7.43, p <0.001 |
| Expertise*Spatial shuffling | DLPFC (-45, 27, 24) | | t(1,35) = 5.87, p <0.001 |
| Expertise*Spatial shuffling | R INS (33, 27, -3) | 943 | t(1,35) = 7.31, p <0.001 |
| Expertise*Spatial shuffling | OTC (-36, -72, 15) | 1740 | t(1,35) = 6.01, p <0.001 |
| Expertise*Spatial shuffling | L IPS (-18, -66, 42) | | t(1,35) = 5.71, p <0.001 |
| Expertise*Spatial shuffling | R IPS (33, -39, 51) | 984 | t(1,35) = 5.89, p <0.001 |
| Expertise*Spatial shuffling | preSMA (9, 21, 42) | 437 | t(1,35) = 4.72, p <0.001 |
| Expertise*Spatial shuffling | R SMA (15, 0, 66) | 295 | t(1,35) = 4.5, p <0.001 |
| Expertise*Spatial shuffling | Vermis 4 5 (9, -54, -24) | 263 | t(1,35) = 4.22, p <0.001 |
| Expertise*Temporal cropping | IPL (-36, -33, 51) | 433 | t(1,35) = 5.19, p = 0.003 |
| Expertise*Temporal cropping | L pMTG (-48, -72, 9) | 138 | t(1,35) = 5.17, p = 0.003 |
| Expertise*Temporal cropping | R pMTG (51, -57, 12) | 190 | t(1,35) = 4.79, p = 0.005 |
| Expertise*Temporal cropping | L DLPFC (-42, 30, 30) | 197 | t(1,35) = 4.6, p = 0.005 |
| Expertise*Temporal cropping | L Post Central Gyrus (-57, -18, 36) | 74 | t(1,35) = 4.5, p = 0.006 |
| Expertise*Temporal cropping | L Cereb 8 (-15, -57, -36) | 23 | t(1,35) = 4.14, p = 0.01 |
| Expertise*Temporal cropping | R IFG (54, 33, 18) | 91 | t(1,35) = 4.1, p = 0.01 |
| Expertise*Temporal cropping | R INS (39, 24, -3) | 32 | t(1,35) = 4.09, p = 0.01 |
| Expertise*Temporal cropping | L SMA (-9, -3, 66) | 15 | t(1,35) = 3.75, p = 0.016 |
| Expertise*Temporal cropping | R preSMA (6, 24, 51) | 27 | t(1,35) = 3.73, p = 0.016 |
| Covariate Expertise*Spatial shuffling | R INS (39, 21, -6) | 741 | t(1,35) = 6.14, p <0.001 |
| Covariate Expertise*Spatial shuffling | R DLPFC (39, 15, 24) | | t(1,35) = 4.94, p <0.001 |
| Covariate Expertise*Spatial shuffling | L INS (-27, 27, -6) | 332 | t(1,35) = 5.97, p <0.001 |
| Covariate Expertise*Spatial shuffling | L DLPFC (-48, 27, 27) | | t(1,35) = 5.23, p <0.001 |
| Covariate Expertise*Spatial shuffling | OTC (-45, -63, -3) | 154 | t(1,35) = 5.03, p <0.001 |
| Covariate Expertise*Spatial shuffling | R IPS (33, -39, 51) | 160 | t(1,35) = 4.78, p = 0.001 |
| Covariate Expertise*Spatial shuffling | L IPS (-27, -69, 30) | 183 | t(1,35) = 4.72, p = 0.001 |
| Covariate Expertise*Spatial shuffling | preSMA (0, 30, 45) | 161 | t(1,35) = 4.56, p = 0.001 |
| Covariate Expertise*Spatial shuffling | L Sup Marg (-63, -30, 27) | 38 | t(1,35) = 4.05, p = 0.001 |
| Covariate Expertise*Spatial shuffling | R Caudate (9, 12, 6) | 33 | t(1,35) = 3.99 p = 0.001 |
| Covariate Expertise*Spatial shuffling | R Temp Mid (39, -60, 9) | 24 | t(1,35) = 3.91 p = 0.001 |
| Covariate Expertise*Spatial shuffling | L Cereb 7b (-9, -72, -42) | 24 | t(1,35) = 3.73 p = 0.001 |
| Covariate Expertise*Temporal cropping | IPL (-36, -36, 48) | 1121 | t(1,35) = 5.56, p = 0.001 |
| Covariate Expertise*Temporal cropping | L pMTG (-48, -75, 9) | 220 | t(1,35) = 4.66, p = 0.003 |
| Covariate Expertise*Temporal cropping | L INS (-30, 27, 0) | 113 | t(1,35) = 4.52, p = 0.004 |
| Covariate Expertise*Temporal cropping | L RO (-45, 6, 15) | 299 | t(1,35) = 4.47, p = 0.004 |
| Covariate Expertise*Temporal cropping | L SMA (-9, -21, 48) | 173 | t(1,35) = 4.31, p = 0.004 |
| Covariate Expertise*Temporal cropping | R IFG pars tri (51, 36, 21) | 39 | t(1,35) = 3.97, p = 0.007 |
| Covariate Expertise*Temporal cropping | R Cereb 6 (21, -48, -27) | 65 | t(1,35) = 3.96, p = 0.007 |
| Covariate Expertise*Temporal cropping | L Cereb 8 (-15, -57, -33) | 29 | t(1,35) = 3.96, p = 0.007 |
| Covariate Expertise*Temporal cropping | R INS (33, 30, -6) | 42 | t(1,35) = 3.86, p = 0.009 |
| Covariate Expertise*Temporal cropping | L Sup Marg (-48, -39, 27) | 32 | t(1,35) = 3.7, p = 0.011 |
| Covariate Expertise*Temporal cropping | R pMTG (42, -63, 3) | 148 | t(1,35) = 3.55, p = 0.015 |

Brain regions showing significant greater activity for the three contrasts: musician > control * piano > forte, musician > control * temporally cropped > unmodified, musician > control * spatially shuffled > unmodified. For all contrasts, significance is corrected at the peak using p<.05 false discovery rate (FDR). Minimal voxel size k = 10.

aITG: inferior temporal gyrus, anterior part; aMTG: medial temporal gyrus, anterior part; aSTG: supperior temporagl gyrus, anterior part; aTFus: temporal fusiform, anterior part; Cereb6: cerebellum 6; Cereb1: cerebellum 1; Cereb3: cerebellum 3; Cereb8: cerebellum 8; DLFC: dorso lateral frontal cortex; DLPFC: dorso lateral prefrontal cortex; FO: frontal operculum; FP: frontal pole; IFG oper: inferior frontal gyrus operculum; IFG pars tri: inferior frontal gyrus pars triangularis; INS: insula; IPL: inferior parietal lobule; IPS: inferior parietal sulcus; L: left; LG: lingual gyrus; MFC: medial frontal cortex; MTG: medial temporal gyrus; OTC: occipito-temporal cortex; PFC: prefrontal cortex; pITG: inferior temporal gyrus, posterior part; pMTG: medial temporal gyrus, posterior part; Post Central Gyrus: posterior central gyrus; Post Cing: posterior cingulate; PostCG: posterior cingulate gyrus; preSMA: pre supplementary motor area; pSMG: superior medial gyrus, posterior part; pTFusC: temporal fusiform cortex, posterior part; R: right; RO: rolandic operculum; SMA: superior motor area; SMG: superior medial gyrus; SPL: superior parietal lobule; SubCalC: subcallosal cortex; SupMarg: supramarginal; toITG: inferior temporal gyrus, temporooccipital part; toMTG: medial temporal gyrus, temporooccipital part; Temp Mid: temporal middle; Ver7: Vernis 7; Ver8: Vernis 8;

#### Supplementary Table 5

**ANOVA Betas for each region of interest**

| Model | Region of interest (MNI mm [x,y,z]) | F-test |
|---|---|---|
| Expertise*Nuance | DLPFC (-42 12 30) | F(1,35) = 3.16, p = 0.08 |
| Expertise*Nuance | preSMA (0 27 45) | F(1,35) = 9.94, p = 0.002 |
| Expertise*Spatial shuffling | DLPFC (-45 27 24) | F(1,35) = 10.92, p = 0.002 |
| Expertise*Spatial shuffling | L INS (-27 27 -3) | F(1,35) = 8.1, p = 0.006 |
| Expertise*Spatial shuffling | L IPS (-18 -66 42) | F(1,35) = 5.32, p = 0.024 |
| Expertise*Spatial shuffling | OTC (-36 -72 15) | F(1,35) = 2.14, p = 0.148 |
| Expertise*Spatial shuffling | preSMA (9 21 42) | F(1,35) = 3.9, p = 0.052 |
| Expertise*Spatial shuffling | R INS (33 27 -3) | F(1,35) = 13.78, p < 0.001 |
| Expertise*Spatial shuffling | R IPS (33 -39 51) | F(1,35) = 3.66, p = 0.06 |
| Expertise*Temporal cropping | IPL (-36 -33 51) | F(1,35) = 2.41, p = 0.125 |
| Expertise*Temporal cropping | L pMTG (-48 -72 9) | F(1,35) = 0.6, p = 0.441 |
| Expertise*Temporal cropping | R pMTG (51 -57 12) | F(1,35) = 1.11, p = 0.295 |
| Expertise*Temporal cropping | R preSMA (6 24 51) | F(1,35) = 2.2, p = 0.143 |
| Expertise*Temporal cropping | L DLPFC (-42 30 30) | F(1,35) = 5.22, p = 0.026 |
| Expertise*Temporal cropping | L Post Central Gyrus (-57 -18 36) | F(1,35) = 1.84, p = 0.18 |
| Expertise*Temporal cropping | L Cereb 8 (-15 -57 -36) | F(1,35) = 2.64, p = 0.109 |
| Expertise*Temporal cropping | R IFG pars tri (54 33 18) | F(1,35) = 2.37, p = 0.128 |
| Expertise*Temporal cropping | R INS (39 24 -3) | F(1,35) = 2.05, p = 0.157 |
| Expertise*Temporal cropping | L SMA (-9 -3 66) | F(1,35) = 1.84, p = 0.179 |

ANOVA comparing the interaction between the expertise and the different conditions for the beta extracted from the SPM.mat of first-level analysis.

aITG: inferior temporal gyrus, anterior part; aMTG: medial temporal gyrus, anterior part; aSTG: supperior temporagl gyrus, anterior part; aTFus: temporal fusiform, anterior part; Cereb6: cerebellum 6; Cereb1: cerebellum 1; Cereb3: cerebellum 3; Cereb8: cerebellum 8; DLFC: dorso lateral frontal cortex; DLPFC: dorso lateral prefrontal cortex; FO: frontal operculum; FP: frontal pole; IFG oper: inferior frontal gyrus operculum; IFG pars tri: inferior frontal gyrus pars triangularis; INS: insula; IPL: inferior parietal lobule; IPS: inferior parietal sulcus; L: left; LG: lingual gyrus; MFC: medial frontal cortex; MTG: medial temporal gyrus; OTC: occipito-temporal cortex; PFC: prefrontal cortex; pITG: inferior temporal gyrus, posterior part; pMTG: medial temporal gyrus, posterior part; Post Central Gyrus: posterior central gyrus; Post Cing: posterior cingulate; PostCG: posterior cingulate gyrus; preSMA: pre supplementary motor area; pSMG: superior medial gyrus, posterior part; pTFusC: temporal fusiform cortex, posterior part; R: right; RO: rolandic operculum; SMA: superior motor area; SMG: superior medial gyrus; SPL: superior parietal lobule; SubCalC: subcallosal cortex; SupMarg: supramarginal; toITG: inferior temporal gyrus, temporooccipital part; toMTG: medial temporal gyrus, temporooccipital part; Ver7: Vernis 7; Ver8: Vernis 8;

#### Supplementary Table 6

**Connectivity Statistics**

| Model | Seed | Target | Statistics |
|---|---|---|---|
| Expertise*Nuance | aTFusC L | aMTG L | T(33) = 4.06, p = 0.043 |
| Expertise*Nuance | Cereb3 L | Cereb10 L | T(33) = 4.60, p = 0.009 |
| Expertise*Nuance | SubCalC | MFC | T(33) = -4.13, p = 0.027 |
| Expertise*Nuance | SubCalC | PostCing | T(33) = -3.98, p = 0.027 |
| Expertise*Nuance | DLPFC L | IPS L | T(33) = -4.19, p = 0.03 |
| Expertise*Nuance | DLPFC L | toITGl | T(33) = -3.89, p = 0.034 |
| Expertise*Nuance | Brain-Stem | LG R | T(33) = 4.42, p = 0.015 |
| Expertise*Nuance | Brain-Stem | LG L | T(33) = 3.82, p = 0.032 |
| Expertise*Nuance | Brain-Stem | Cereb8 L | T(33) = 3.78, p = 0.032 |
| Expertise*Nuance | MFC | SubCalC | T(33) = -4.13, p = 0.035 |
| Expertise*Nuance | Putamen L | aMTG R | T(33) = 4.19, p = 0.029 |
| Expertise*Nuance | FP L | FO R | T(33) = -4.01, p = 0.05 |
| Expertise*Nuance | aMTG R | Putamen L | T(33) = 4.19, p = 0.029 |
| Expertise*Nuance | LG R | Brain-Stem | T(33) = 4.42, p = 0.015 |
| Expertise*Nuance | aMTG L | aTFusC L | T(33) = 4.06, p = 0.043 |
| Expertise*Nuance | aMTG L | pTFusC R | T(33) = 3.77, p = 0.048 |
| Expertise*Nuance | IPS L | DLPFC L | T(33) = -4.19, p = 0.03 |
| Expertise*Nuance | Cereb10 L | Cereb3 L | T(33) = 4.60, p = 0.009 |
| Expertise*Nuance | FO R | FP L | T(33) = -4.01, p = 0.05 |
| Expertise*Spatial shuffling | IFG oper L | PostCG R | T(33) = -4.50, p = 0.012 |
| Expertise*Spatial shuffling | aITG L | pMTG L | T(33) = 3.81, p = 0.038 |
| Expertise*Spatial shuffling | aITG L | aMTG L | T(33) = 3.75, p = 0.038 |
| Expertise*Spatial shuffling | aITG L | SMG R | T(33) = 3.71, p = 0.038 |
| Expertise*Spatial shuffling | aITG L | aSTG L | T(33) = 3.53, p = 0.048 |
| Expertise*Spatial shuffling | pITG R | Brain-Stem | T(33) = -4.30, p = 0.021 |
| Expertise*Spatial shuffling | Brain-Stem | pITG R | T(33) = -4.30, p = 0.021 |
| Expertise*Spatial shuffling | PostCG R | IFG oper L | T(33) = -4.50, p = 0.012 |
| Expertise*Temporal cropping | Ver8 | IFG oper L | T(33) = 5.31, p = 0.001 |
| Expertise*Temporal cropping | Ver8 | DLPFC | T(33) = 4.58, p = 0.004 |
| Expertise*Temporal cropping | Ver8 | toMTGl | T(33) = 4.52, p = 0.004 |
| Expertise*Temporal cropping | Ver8 | pSMGl | T(33) = 4.02, p = 0.012 |
| Expertise*Temporal cropping | Ver8 | IPS R | T(33) = 3.51, p = 0.037 |
| Expertise*Temporal cropping | Ver8 | IFG oper R | T(33) = 3.47, p = 0.037 |
| Expertise*Temporal cropping | Ver8 | INS L | T(33) = 3.35, p = 0.045 |
| Expertise*Temporal cropping | Ver8 | DLPFC L | T(33) = 3.29, p = 0.045 |
| Expertise*Temporal cropping | SPLl | Ver7 | T(33) = 4.04, p = 0.046 |
| Expertise*Temporal cropping | IFG oper L | Ver8 | T(33) = 5.31, p = 0.001 |
| Expertise*Temporal cropping | to MTG L | Ver8 | T(33) = 4.52, p = 0.012 |
| Expertise*Temporal cropping | Ver7 | SPL L | T(33) = 4.04, p = 0.046 |
| Expertise*Temporal cropping | DLPFC | Ver8 | T(33) = 4.58, p = 0.01 |
| Expertise*Temporal cropping | pSMGl | Ver8 | T(33) = 4.02, p = 0.048 |

Statistics associated with the different connections in the connectivity analysis.
P value is FDR corrected at p<.05; aITG: inferior temporal gyrus, anterior part; aMTG: medial temporal gyrus, anterior part; aSTG: supperior temporagl gyrus, anterior part; aTFus: temporal fusiform, anterior part; Cereb6: cerebellum 6; Cereb1: cerebellum 1; Cereb3: cerebellum 3; Cereb8: cerebellum 8; DLFC: dorso lateral frontal cortex; DLPFC: dorso lateral prefrontal cortex; FO: frontal operculum; FP: frontal pole; IFG oper: inferior frontal gyrus operculum; IFG pars tri: inferior frontal gyrus pars triangularis; INS: insula; IPL: inferior parietal lobule; IPS: inferior parietal sulcus; L: left; LG: lingual gyrus; MFC: medial frontal cortex; MTG: medial temporal gyrus; OTC: occipito-temporal cortex; PFC: prefrontal cortex; pITG: inferior temporal gyrus, posterior part; pMTG: medial temporal gyrus, posterior part; Post Cing: posterior cingulate; PostCG: posterior central gyrus; preSMA: pre supplementary motor area; pSMG: superior medial gyrus, posterior part; pTFusC: temporal fusiform cortex, posterior part; R: right; RO: rolandic operculum; SMA: superior motor area; SMG: superior medial gyrus; SPL: superior parietal lobule; SubCalC: subcallosal cortex; SupMarg: supramarginal; toITG: inferior temporal gyrus, temporooccipital part; toMTG: medial temporal gyrus, temporooccipital part; Ver7: Vernis 7; Ver8: Vernis 8;

**Supplementary Figure 1**

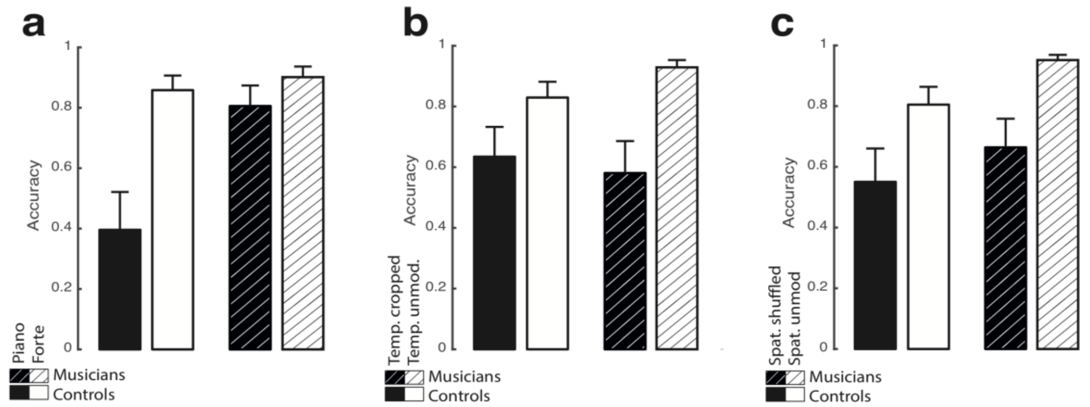


Behavioral performance of both groups for every interaction when the sex is captured by random intercept effect. (**a**) Averaged performance per group for piano vs. forte piece dynamics. (**b**) Averaged performance per group for piano vs. forte piece dynamics in temporally cropped vs. temporally unmodified sequences. (**c**) Averaged performance per group for piano vs. forte piece dynamics in spatially shuffled vs. spatially unmodified sequences. The present results highlight the behavioral performance of the participants when the variance associated with sex is captured by the model. Similar patterns can be observed with or without sex being captured by random effects.

**Supplementary Figure 2**

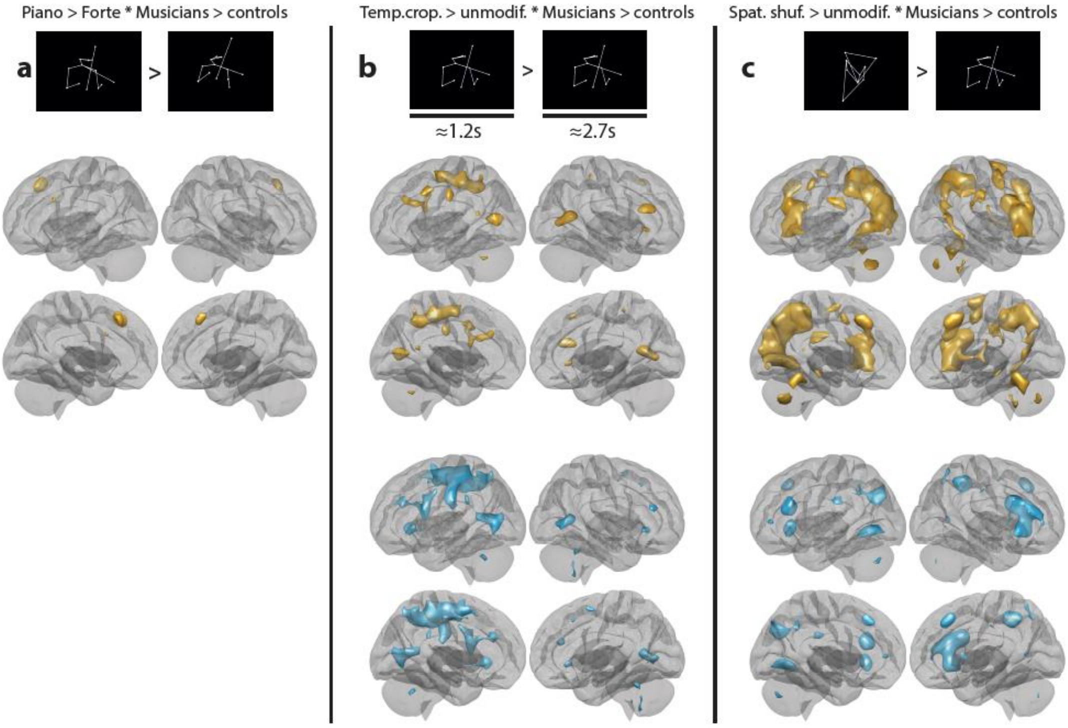


Activations represented in volumes in the brain for the different conditions. (a) Piano>Forte * Musicians > Controls. (b) Temporally cropped>Temporally unmodified * Musicians > Controls. (c) Spatially shuffled>Spatially unmodified * Musicians > Controls. Gold clusters are the effect of condition contrasts while teal clusters reflect the interaction with the group-level general task performance covariate. $p<.05$, FDR corrected at the peak level.

#### Supplementary Figure 3

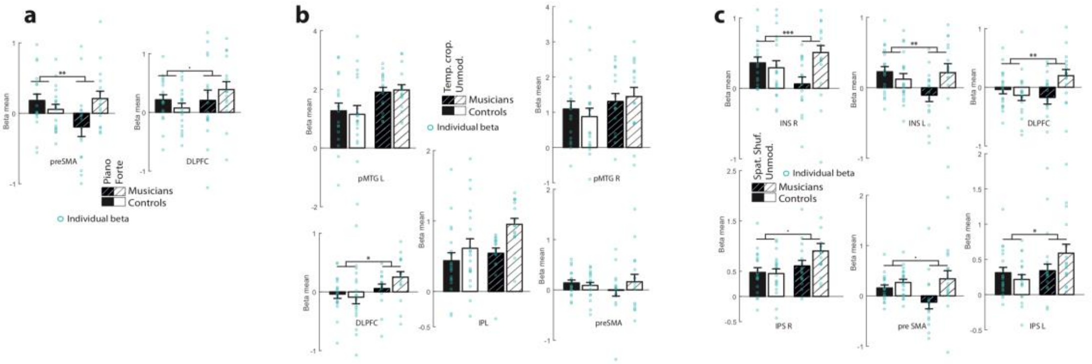


(**a**) Beta values of the general linear model (GLM) for piano and forte sequences for musicians and control participants. (**b**) Beta values of the GLM for temporally cropped and unmodified sequences for musicians and control participants. (**c**) Beta values of the GLM for spatially shuffled and unmodified sequences for musicians and control participants.

aITG: inferior temporal gyrus, anterior part; aMTG: medial temporal gyrus, anterior part; aSTG: supperior temporagl gyrus, anterior part; aTFus: temporal fusiform, anterior part; Cereb6: cerebellum 6; Cereb1: cerebellum 1; Cereb3: cerebellum 3; Cereb8: cerebellum 8; DLFC: dorso lateral frontal cortex; DLPFC: dorso lateral prefrontal cortex; FO: frontal operculum; FP: frontal pole; IFG oper: inferior frontal gyrus operculum; IFG pars tri: inferior frontal gyrus pars triangularis; INS: insula; IPL: inferior parietal lobule; IPS: inferior parietal sulcus; L: left; LG: lingual gyrus; MFC: medial frontal cortex; MTG: medial temporal gyrus; OTC: occipito-temporal cortex; PFC: prefrontal cortex; pITG: inferior temporal gyrus, posterior part; pMTG: medial temporal gyrus, posterior part; Post Central Gyrus: posterior central gyrus; Post Cing: posterior cingulate; PostCG: posterior cingulate gyrus; preSMA: pre supplementary motor area; pSMG: superior medial gyrus, posterior part; pTFusC: temporal fusiform cortex, posterior part; R: right; RO: rolandic operculum; SMA: superior motor area; SMG: superior medial gyrus; SPL: superior parietal lobule; SubCalC: subcallosal cortex; SupMarg: supramarginal; toITG: inferior temporal gyrus, temporooccipital part; toMTG: medial temporal gyrus, temporooccipital part; Ver7: Vernis 7; Ver8: Vernis 8;

**Supplementary Figure 4**

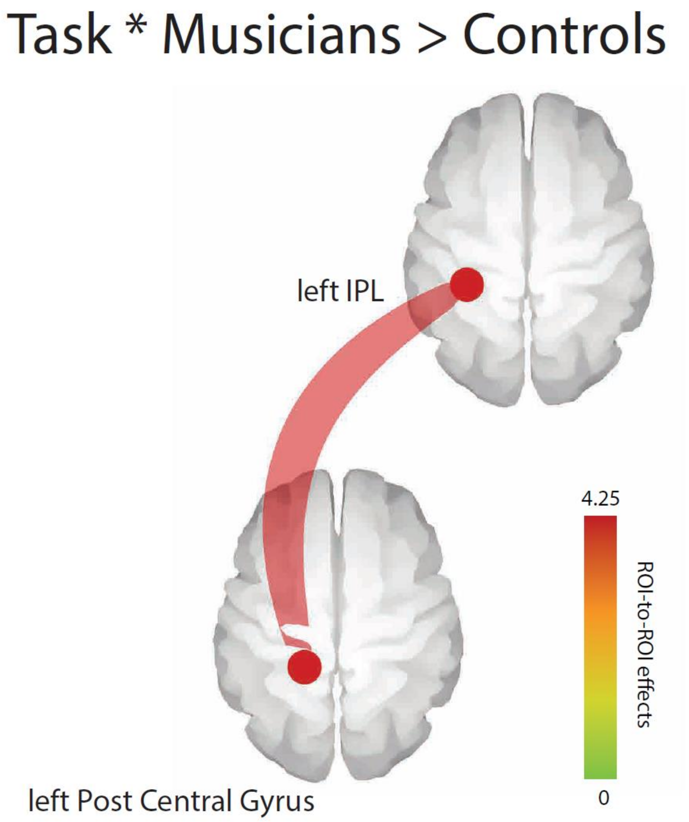


Functional connectivity between regions of interest highlighted in the GLM analysis for the main effect of the task.
